# Evolutionary branching of male emergence timing: Trade-offs and variance asymmetry as drivers of dimorphism

**DOI:** 10.64898/2026.04.10.717617

**Authors:** Hidaka Kubo, Ryo Yamaguchi, Yuuya Tachiki

**Affiliations:** Graduate school of Life Science, Hokkaido University, Sapporo, Hokkaido, Japan; Department of Advanced Transdisciplinary Science, Hokkaido University, Sapporo, Hokkaido, Japan; Department of Biological Sciences, Tokyo Metropolitan University, Hachioji, Tokyo, Japan

**Author notes:** Corresponding author: Hidaka Kubo.

## Abstract

Classical models of protandry predict unimodal male emergence timing, yet empirical observations in butterflies and bees reveal dimorphism: early-emerging small males coexist with late-emerging large males. The evolutionary mechanisms underlying such discrete alternative reproductive strategies in emergence timing remain poorly understood. In this study, we developed a mathematical model using an adaptive dynamics framework to investigate the conditions under which dimorphic male emergence timing evolves. We explored two potential mechanisms: (1) a trade-off between emergence timing and male competitiveness, and (2) differences in the variance of emergence timing between the sexes. Our analysis demonstrates that both mechanisms produce evolutionary branching, and that extreme variance asymmetry between sexes can generate multiple branching events, yielding three or more distinct male clusters. These findings provide a theoretical foundation for understanding the evolution of alternative reproductive strategies in emergence timing, with implications for other systems where reproductive success depends on temporal overlap between the sexes. We provide testable predictions that a positive correlation between emergence timing and male body size or competitiveness should be observed under the trade-off mechanism, and that the variance in emergence timing within each male morph should be smaller than that of females under the variance asymmetry mechanism.

## Introduction

Life history strategy indicates the schedule of lifetime events—such as growth, development, and reproduction—that organisms have evolved to maximize fitness (Stearns, 1976; Stott et al., 2024). Central to life history theory is the principle that organisms must allocate finite resources among competing traits, and natural selection optimizes this allocation under the ecological conditions each species faces (Williams, 1966; Stearns, 1989). The optimal solution for this allocation often differs between the sexes, since female reproductive success is generally limited by resource access, while male success is limited by the number of mates (Bateman, 1948; Andersson & Iwasa, 1996). Such divergent selection pressures drive the evolution of sexual dimorphism, which is observed in a wide range of traits (Hedrick & Temeles, 1989).

Within this framework, the timing of reproduction emerges as a critical life history trait, particularly in seasonal environments where breeding opportunities are temporally constrained. Intriguingly, individuals of the same sex within a population sometimes adopt strikingly different, genetically determined reproductive strategies. Such alternative reproductive strategies have fascinated evolutionary biologists because they raise fundamental questions about the maintenance of phenotypic diversity under natural selection (Gross, 1996; Oliveira et al., 2008). Classic examples include three distinct male morphs—independent, satellite, and faeder—in the ruff (*Philomachus pugnax*) (Widemo, 1998; Jukema & Piersma, 2006), and the alpha, beta, and gamma males in the marine isopod (*Paracerceis sculpta*) (Shuster, 1987; Shuster, 1989). These discrete morphs are governed by specific genetic loci (e.g., supergenes or Mendelian alleles) and maintained by frequency-dependent selection (Küpper et al., 2016; Shuster & Wade, 1991). While these spatial and behavioral alternative reproductive strategies have received extensive theoretical and empirical attention (Oliveira et al., 2008; Dougherty et al., 2022), the possibility that discrete alternative morphs in emergence timing might evolve within male populations has been largely unexplored theoretically.

A widespread form of such sex-differential reproductive timing is protandry, where males arrive or mature earlier than females (Morbey & Ydenberg, 2001). Protandry is prevalent in the adult emergence of seasonal insects like butterflies. (Wiklund & Fagerström, 1977). The prevailing explanation for this pattern is the “mate opportunity hypothesis”, suggesting that early emergence of males maximizes their reproductive success by ensuring the opportunity to mate with the females, particularly when females mate only once (Wiklund & Fagerström, 1977; Bulmer 1983; Iwasa et al., 1983; Parker & Courtney, 1983). Alternatively, though not mutually exclusive, it is argued that protandry is a byproduct of sexual size dimorphism, which evolves because fecundity selection favors larger females that require longer development times (Wiklund & Solbreck, 1982; Morbey & Ydenberg, 2001). These selective pressures likely act in concert, jointly shaping the degree of protandry observed in natural populations (Morbey, 2013).

Theoretical frameworks describing the evolution of protandry have been extensively developed to formalize the mate opportunity hypothesis. Several early mathematical models demonstrated that protandry evolves due to sexual differences in mating systems, specifically where males are polygynous and females are monandrous (Wiklund & Fagerström, 1977; Bulmer, 1983; Iwasa et al., 1983; Parker & Courtney, 1983). Subsequent extensions have incorporated stochastic environments, female mortality, female polyandry, and trade-offs between male competitiveness and emergence timing (Iwasa & Haccou, 1994; Zonneveld & Metz, 1991; Zonneveld, 1992, 1996). A common feature of these models, however, is that they predict or implicitly assume unimodal emergence distributions. Males are expected to converge on a single optimal emergence time that balances the benefits of early arrival against the costs of reduced mating opportunities later in the season.

Empirical observations challenge the unimodal emergence distributions expected by these classical theories. In the butterfly *Fabriciana nerippe*, male emergence timing exhibits dimorphism associated with body size: early-emerging (protandrous) males are smaller than males emerging later (Kubo et al., 2025). A similar pattern occurs in the solitary bee *Amegilla dawsoni* (Alcock, 1997), where small males emerge early and large males emerge late. These cases suggest that dimorphism in male emergence timing may represent alternative reproductive strategies, with early males gaining an advantage by emerging early, while late males acquire higher competitive ability through prolonged growth. Yet existing theory cannot explain how such dimorphism evolves and is maintained.

Two ecological factors may generate disruptive selection favoring dimorphic male emergence. First, a trade-off between emergence timing and male competitiveness could maintain two distinct strategies: early males gain by accessing virgin females before competitors arrive, while late males compensate by achieving larger body sizes that enhance their success in scramble competition. The dimorphism can evolve when the fitness of these alternative strategies is balanced. Second, differences in the variance of emergence timing between sexes could drive the evolution of dimorphism. If male emergence is more synchronized than female emergence, virgin females at the peak of male emergence become rapidly depleted, reducing the expected mating success of males with intermediate timing. This creates a fitness landscape with peaks at both tails of the male distribution—analogous to evolutionary branching driven by resource competition (e.g., Doebeli & Dieckmann, 2000), where females represent the resource and males represent the consumers.

In this study, we explored the conditions under which dimorphic male emergence timing can evolve, applying an adaptive dynamics framework (Geritz et al. 1998). In this framework, we assume that emergence timing is a genetically determined trait; thus, the distinct phenotypes correspond to genetically fixed alternative strategies rather than condition-dependent tactics. Specifically, we focus on the trade-off between male competitiveness and emergence timing and on sex differences in the variance of emergence distribution. Our analysis reveals that male dimorphism can arise as an evolutionary consequence of disruptive selection in competition for females. Previous theoretical models assumed restrictive functional forms for the competitiveness-timing trade-offs (Zonneveld, 1996) and typically neglected variance differences in the emergence distributions between sexes. By relaxing these assumptions, we demonstrate that evolutionary branching, and hence the origin of discrete male strategies, occurs under biologically plausible conditions. Our results provide a theoretical foundation for understanding the diversity of reproductive timing strategies in insects and other seasonal breeders, linking the evolution of protandry to the broader phenomenon of alternative reproductive strategies.

## Model and Analysis

### Model

#### • Overview and assumptions

We developed a model to investigate the evolution of male emergence timing using the adaptive dynamics framework (Geritz et al., 1998). Unlike previous models that predict unimodal emergence distributions, our approach allows for evolutionary branching—the evolutionary divergence of a single population into distinct phenotypic clusters. This framework enables us to identify conditions under which dimorphic male emergence timing can evolve as a consequence of disruptive selection. Our model builds on the foundational assumptions of Wiklund and Fagerström (1977): (i) Generations do not overlap; (ii) male and female emergence timing are under genetic control and follow normal distributions; (iii) females mate only once in their lifetime, while males can mate multiple times; (iv) all females become fertilized immediately after their emergence; (v) males have an equal and constant death rate; and (vi) The total population size and sex ratio remain constant across generations. The symbols used in this study are summarized in Table 1.

**Table 1.** Notation and parameters used in this study. Default values are given in square brackets. “*” represents parameters used only in individual-based simulations.

| Parameter | Definition | Value range |
| --- | --- | --- |
| $\tau$ | Mean value of male emergence distribution | Evolutionary variable |
| $\sigma_M$ | Standard deviation of male emergence distribution | 0.5–2.0 [1, 0.5] |
| $\sigma_F$ | Standard deviation of female emergence distribution | 0.5–2.0 [1] |
| $\mu$ | Death rate of adult males | 0.2–0.8 [0.5] |
| $s$ | Inflection point of the competitiveness function | -1–1 [0, 0.3] |
| $h$ | Upper asymptote of the competitiveness function | 1–10 [1, 10] |
| $k$ | Steepness of the competitive function | 1–10 [1, 10] |
| $N_{M\tau}$ | Total number of resident male emergences* | [1000] |
| $N_F$ | Total number of female emergences* | [1000] |
| $u$ | Mutation rate* | [0.05] |
| $\sigma_{mut}$ | Standard deviation of the distribution of mutational effects* | [0.02] |

#### • Probability distribution of emergence timing

We assume that the probability distribution of male and female emergence timing follows a normal distribution with τ and 0 as their respective means. Thus, the probability distribution of male emergence timing *P*_*M*_ and that of female emergence timing *P*_*F*_ are as follows:

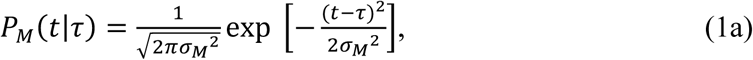

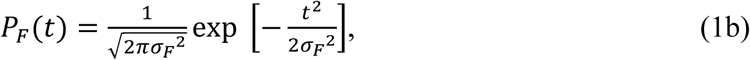

where σ_*M*_ and σ_*F*_ denote standard deviation of *P*_*M*_ and *P*_*F*_ respectively. *P*_*M*_(*t*|τ) represents the probability that male with trait τ emerges at time *t*.

#### • Male competitiveness

In many insects, male body size positively correlates with mating success (Thornhill & Alcock, 1983). Because body size increases with development time, males that emerge later may have higher competitive ability. We capture this relationship through a competitiveness function, *f*(*t*), determined by its emergence time *t*. Males acquire mates in proportion to their competitiveness relative to all competing males (e.g., a male with *f* = 2 obtains twice as many mates as one with *f* = 1 at a given time). At a given time, males with competitiveness *f*(*t*) emerge with probability *P*_*M*_(*t*|τ). We define the “total male competitiveness of males with trait τ at time t” as *N*_*M*_τ*C*(*t*|τ), where *N*_*M*_τ and *C*(*t*|τ) denote the number of males with trait τ, and the competitiveness density of males with trait τ at time t, respectively. Therefore, the competitiveness density of males increases at a rate of *f*(*t*)*P*_*M*_(*t*|τ). Conversely, the competitiveness density decreases due to the death rate, μ. Thus, *C*(*t*|τ) satisfies the following differential equation:

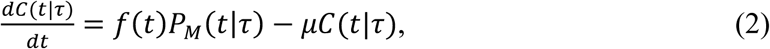

Under the initial condition 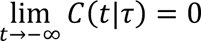, the solution of Eq. (2) is:

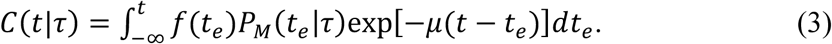

where, *t*_*e*_ denotes the emergence time of a male.

#### • Invasion fitness

We use the adaptive dynamics framework to study the long-term evolutionary dynamics of male emergence timing. We introduce a rare mutant τ′ attempting to invade a monomorphic resident trait τ. We assume that the magnitude of mutation |τ^′^ − τ| is infinitesimally small. To assess the possibility of mutant invasion, we define the invasion fitness *W*, which represents the growth rate of a single mutant introduced into a resident population. In this model, the invasion fitness is defined as the lifetime number of matings per mutant male.

Under assumptions (iii) and (iv), all emerging females mate immediately only once and become fertilized. The number of females emerging at time *t* is given by *N*_*F*_*P*_*F*_(*t*), where *N*_*F*_ denotes the total number of female emergence. A male that emerged at time *t*_*e*_ obtains matings in proportion to his competitiveness *f*(*t*_*e*_) relative to the total competitiveness of all males present. When both resident (τ) and mutant (τ′) males are present, the total competitiveness at time *t* is given by *N*_*M*_τ*C*(*t*|τ) + *N*_*M*_ ′ *C*(*t*|τ^′^), where *N*_*M*_τ and *N*_*M*_τ′ denote the total numbers of resident and mutant males, respectively. Thus, the mating rate of a male with competitiveness *f*(*t*_*e*_) at time *t*, *M*(*t*_*e*_, *t*|τ, τ′) can be written as:

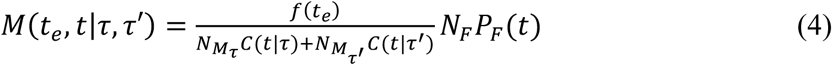

The total number of matings for a male that emerge at time *t*_*e*_, denoted by ϕ, can be obtained by integrating the product of *M*(*t*_*e*_, *t*|τ, τ′) and survival probability from *t*_*e*_:

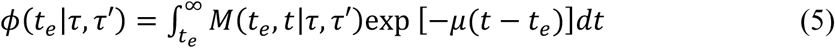

Thus, the total number of matings for a mutant with trait τ′, denoted by *W*(τ^′^|τ), can be obtained by calculating the expected value of ϕ using *P*_*M*_(*t*_*e*_|τ′):

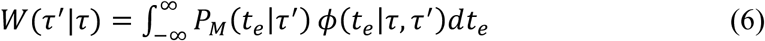

Taking the limit as *N*_*M*_τ′ → 0, and assuming *N*_*M*_τ = *N*_*F*_, we obtain the following:

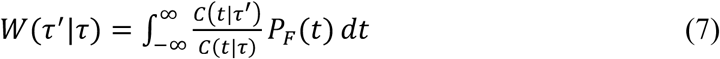

#### • Competitiveness function

In this study, we use the following sigmoid function as the male competitiveness function.

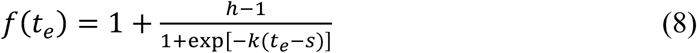

where *s*, *h* and *k* denote the inflection point, upper asymptote, and steepness of the competitive function, respectively. This sigmoid form captures the biological reality that male body size (and thus competitiveness) increases gradually and monotonically with development time, eventually reaching an asymptotic maximum due to physiological constraints. The shape of this function is shown in Fig. S1.

## Model Analysis

### • Evolutionarily singular strategy

To investigate the evolutionary direction of male emergence timing, we calculate the local fitness gradient *D*(τ) (Geritz et al., 1998).

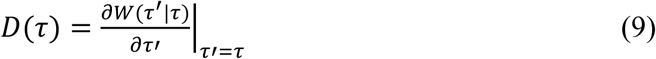

The sign of the local fitness gradient determines the direction of selection acting on the emergence timing. A positive gradient indicates that selection favors a mutant with later emergence (larger τ), whereas a negative gradient favors a mutant with earlier emergence (smaller τ). By solving *D*(τ) = 0, we obtain an evolutionary equilibrium τ^∗^ at which selection ceases, referred to as the evolutionarily singular strategy.

### • Stability of singular strategy

A singular strategy is “convergence stable” (Christiansen 1991) if a resident population near τ^∗^ can be invaded by mutants that are even closer to τ^∗^. Around a convergence stable singular strategy τ^∗^, the sign of the local fitness gradient changes from positive to negative, which means that *D*(τ) is a (locally) decreasing function of τ. Hence, the condition for convergence stability is given as follows:

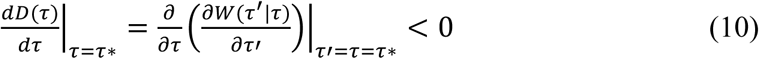

A singular strategy is “evolutionarily stable strategy” (ESS) (Maynard Smith, 1974) if no mutant can invade in a neighborhood of τ^∗^ which implies that the second derivative of invasion fitness evaluated at the singular strategy is negative:

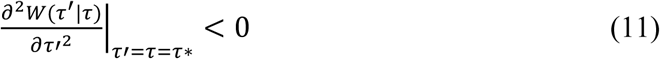

The condition in which a singular strategy τ^∗^ is convergence stable but not evolutionarily stable indicates that an initially monomorphic population is expected to undergo evolutionary branching toward a dimorphic population.

### • Individual-based simulation

We conducted an individual-based simulation of the evolutionary dynamics. The numbers of male and female individuals in each generation are fixed at *N*_*M*_τ and *N*_*F*_, respectively. The probability of producing offspring of a focal male is proportional to the male’s fitness, and offspring inherit their father’s trait value. Mutations occur with mutation rate *u* and are chosen randomly from a Gaussian distribution with the mean 0 and the standard deviation σ_*mut*_. The simulation codes are available on GitHub (https://github.com/hidakakubo/AdaptiveDynamics_protandry).

## Result

We analyzed the evolutionary dynamics of male emergence timing under three conditions: (1) Neutral: no trade-off between emergence timing and competitiveness (*c*(*t*) = 1), with equal variance in emergence timing for both sexes (σ_*M*_ = σ_*F*_); (2) Trade-off: a trade-off exists between emergence timing and competitiveness ( *c*(*t*) = 1 +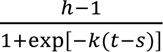 and σ_*M*_ = σ_*F*_); and (3) Different-variance: males and females have different variances in emergence timing (*c*(*t*) = 1 and σ_*M*_ ≠ σ_*F*_). Our results reveal that evolutionary branching can occur under the trade-off and different-variance conditions, but not under the neutral condition. We then examined how the occurrence of evolutionary branching depends on various biological parameter values. Furthermore, our results demonstrate that multiple branching events, resulting in the evolution of three or more distinct trait clusters, can occur under the different-variance condition, particularly when the variance in male emergence timing is substantially smaller than that of females.

### Evolutionary outcome under different conditions

To investigate the evolutionary outcome, we constructed pairwise invasibility plots (PIPs) (Geritz et al. 1998) for the neutral (Fig. S2(a)), trade-off (Fig. 2(a, d)), and different- variance (Fig. 3(a, d)) conditions. PIPs provide an intuitive graphical tool for analyzing the convergence and evolutionary stability of singular strategies. In PIPs, invasion fitness *W*(τ^′^|τ) is plotted for all combinations of resident trait values τ (horizontal) and mutant trait values τ′ (vertical).

**Fig. 1.**
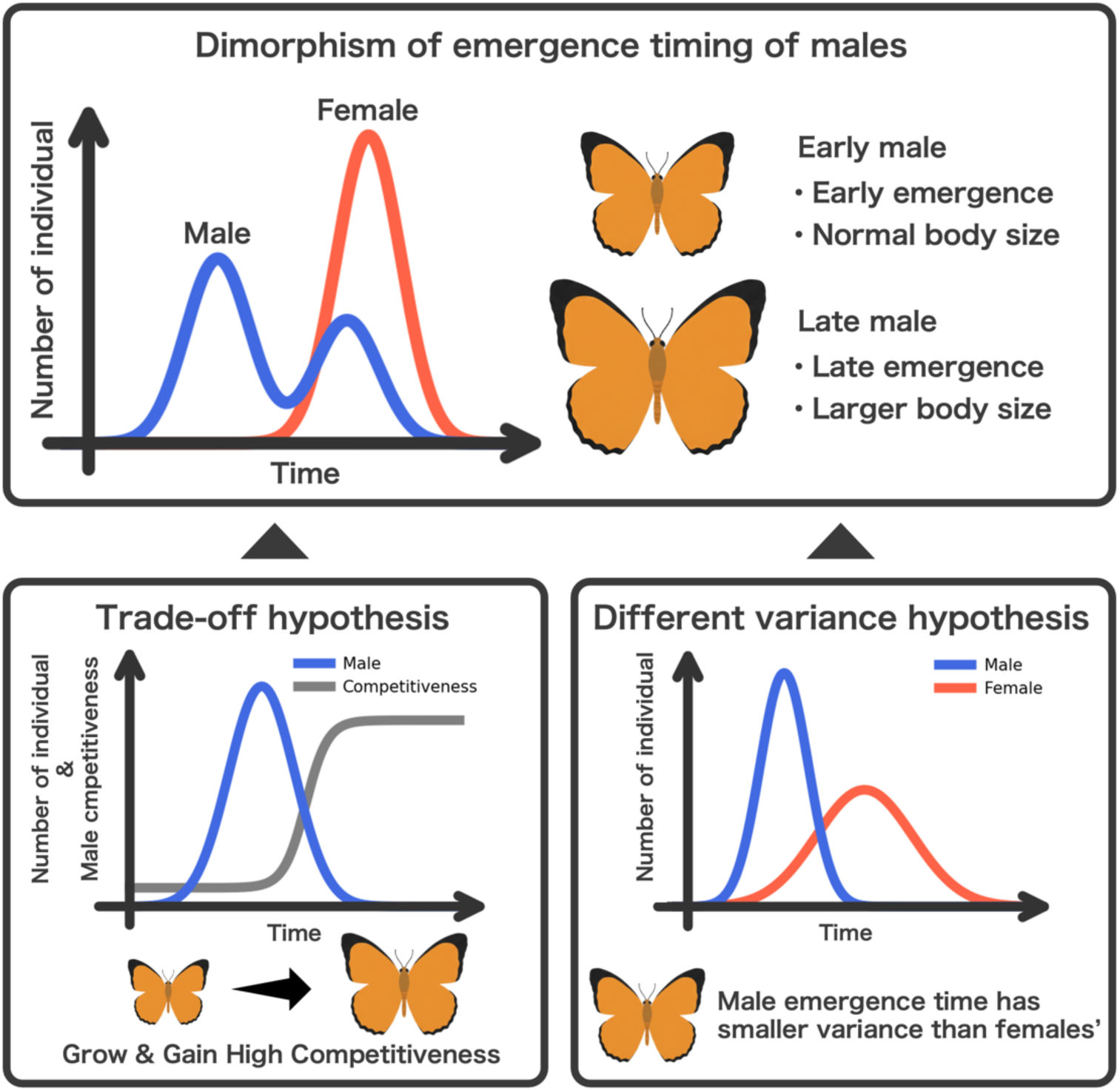
Schematic illustration of the dimorphism of male emergence timing and the two alternative hypotheses for its evolutionary factor of it. The top panel shows the dimorphism in emergence timing, which is the focus of this study. The vertical axis represents the number of individuals, and the horizontal axis represents time in a reproductive season. The distribution of male individuals is bimodal. Males that emerge earlier are called Early males (with normal body size), and those that emerge later are called Late males (with larger body size). The bottom left panel presents the trade-off hypothesis. This hypothesis proposes that late-emerging males grow and gain a larger body size, which provides them with a highly competitive advantage in the competition for females. In this situation, the benefits of early emergence and late emergence are balanced, leading to the evolution of the dimorphism. The bottom right panel presents the different-variance hypothesis. This hypothesis posits that the variance in male emergence timing is smaller than that of females. In this situation, the concentrated emergence of males leads to a high consumption of females at a specific time, which reduces the fitness of males with intermediate traits, causing disruptive selection.

**Fig. 2.**
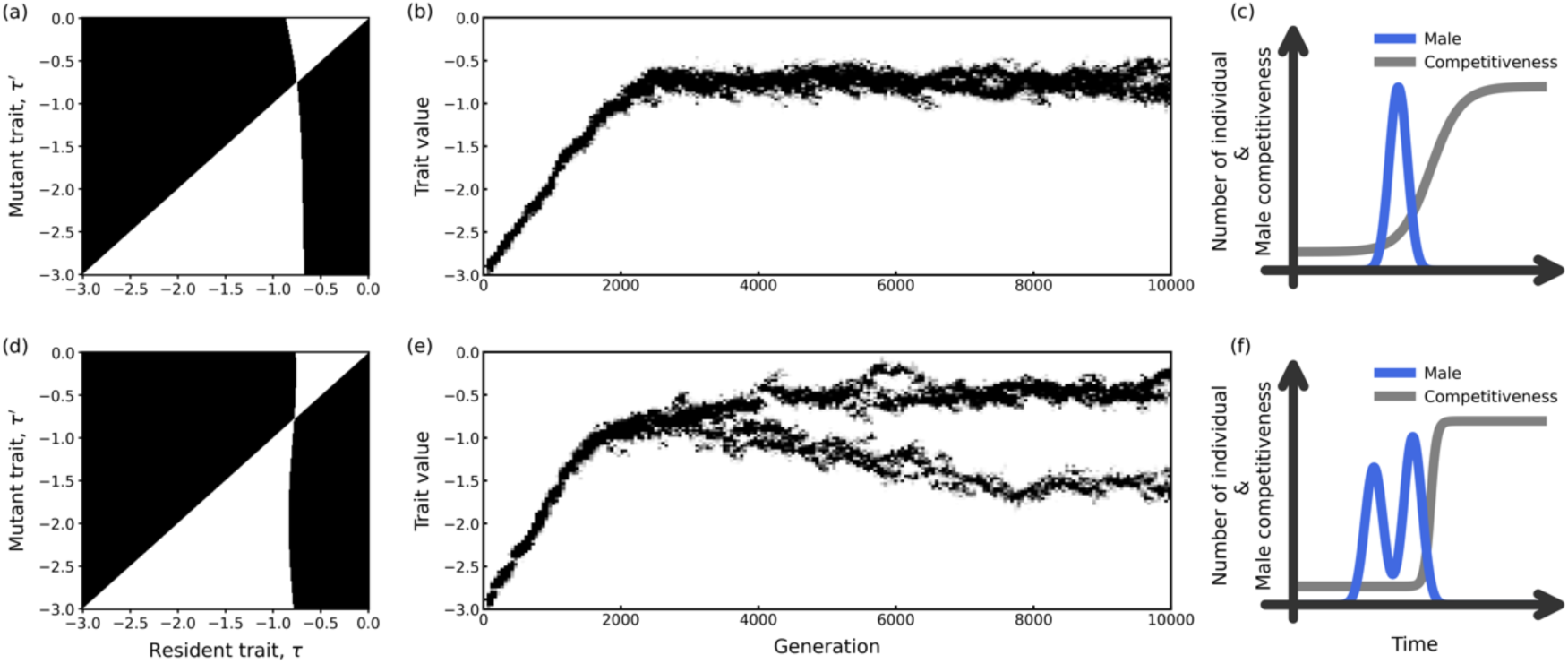
Pairwise invasibility plots (PIPs) and evolutionary dynamics generated by the individual-based simulation under the trade-off conditions. (a, b, c) Branching condition with *k* = 2.0. (d, e, f).Non-branching condition with *k* = 10.0. (a, d) PIPs showing invasion fitness *W*(τ′|τ) where τ and τ′ represent resident and mutant trait values, respectively. Black regions indicate successful invasion (*W* > 1). (b), (e) Individual-based simulation results showing trait distribution over 10000 generations. (c), (f) Schematic plot of each evolutionary outcome. Parameters are σ_*M*_ = 1.0, σ_*F*_ = 1.0, μ = 0.5, *s* = 0.3, *h* = 10.0, *N*_*M*_τ = 1000, *N*_*F*_ = 1000, *u* = 0.05, and σ_*mut*_ = 0.02.

**Fig. 3.**
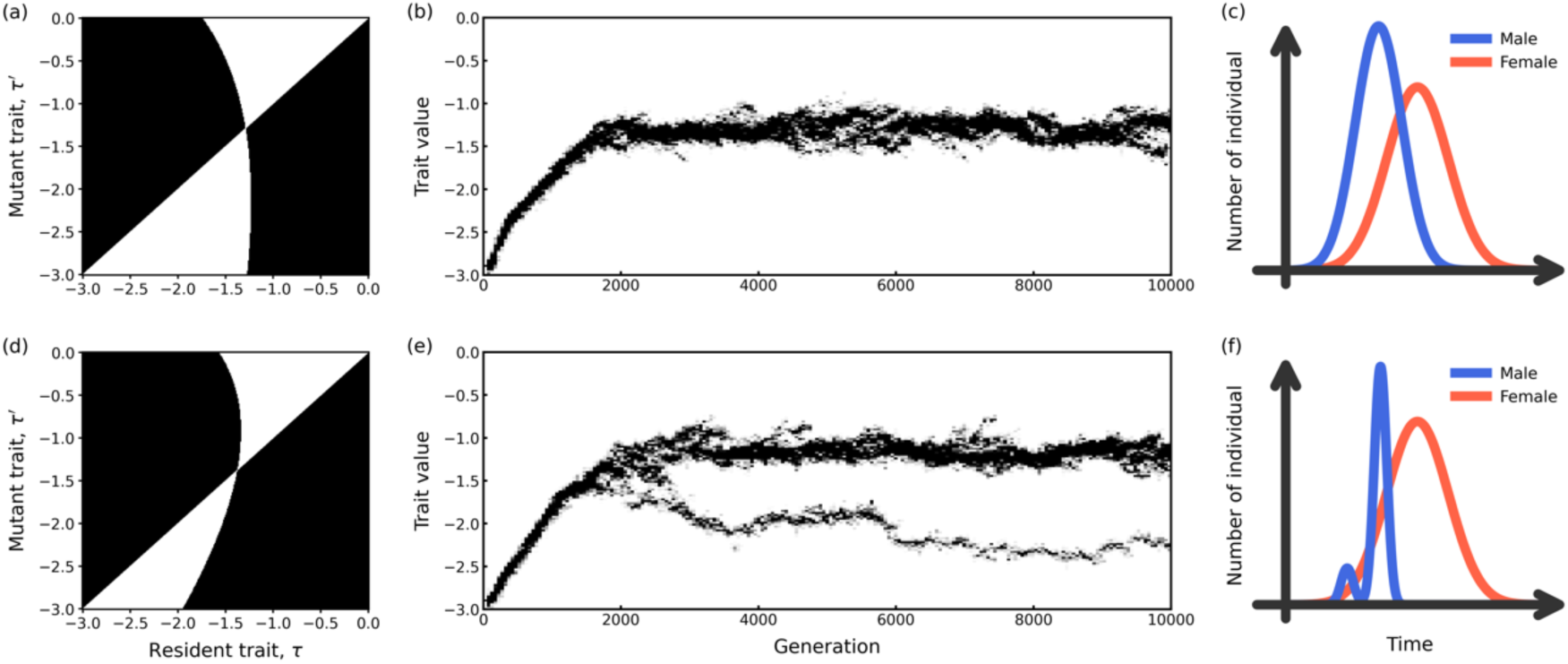
Pairwise invasibility plots (PIPs) and evolutionary dynamics generated by the individual-based simulation under the different-variance conditions. (a, b, c) Branching condition with σ_*M*_ = 0.75. (d, e, f) Non-branching condition with σ_*M*_ = 0.5. (a, d) PIPs showing invasion fitness *W*(τ′|τ) where τ and τ′ represent resident and mutant trait values, respectively. Black regions indicate successful invasion ( *W* > 1). (b), (e) Individual-based simulation results showing trait distribution over 10000 generations. (c),(f) Schematic plot of each evolutionary outcome. Parameters are σ_*F*_ = 1.0, μ = 0.5, *s* = 0.0, *h* = 1.0, *k* = 1.0, *N*_*M*_τ = 1000, *N*_*F*_ = 1000, *u* = 0.05, and σ_*mut*_ = 0.02.

Black regions indicate *W*(τ^′^|τ) > 1 (successful mutant invasion), while white regions indicate *W*(τ^′^|τ) < 1 (unsuccessful invasion). The diagonal line indicates neutral mutants, where the mutant strategy is identical to that of the resident (i.e., *W*(τ|τ) = 1). The intersection of the diagonal line and another line satisfying *W*(τ^′^|τ) = 1 corresponds to the evolutionarily singular strategy, where the local fitness gradient (eq. 9) equals zero. It was confirmed that there was only one singular strategy in the obtained results.

A singular strategy is convergence stable if, in its vicinity, mutants with trait values closer to the singular strategy can invade—graphically, this corresponds to black regions above the diagonal to the left of the singular strategy and below the diagonal to its right. Accordingly, Fig. S2(a), Fig. 2(a, d), and Fig. 3(a, d) are all convergence stable, respectively. In addition, a singular strategy is evolutionarily stable (an ESS) if the vertical line passing through it lies entirely within the white region, indicating that no nearby mutants can invade (i.e., Fig. S2(a), Fig. 2(a), and Fig. 3(a)). In contrast, if this vertical line intersects black regions (i.e., Fig. 2(d), and Fig. 3(d)), the singular strategy can be invaded by both larger and smaller trait values. Such singular strategies are evolutionary branching points, at which disruptive selection drives the evolution of polymorphism.

Under the neutral condition, the singular strategy is both convergence and evolutionarily stable (Fig. S2(a)). Individual-based simulations confirmed that the trait values converged to the singular point predicted by the PIP and remained stable thereafter (Fig. S2(b)), regardless of initial trait values. Notably, the ESS trait value is negative, indicating that male emergence timing exhibits protandry.

Under the trade-off condition, the singular strategy is always convergence stable; however, its evolutionary stability depends on parameter values. When the steepness of the competitiveness function *k* is small, the singular strategy is an ESS (Fig. 2(a)), and individual-based simulations confirmed convergence to a stable monomorphic population (Fig. 2(b)). Conversely, when *k* is large, the singular strategy becomes an evolutionary branching point (Fig. 2(d)). In this case, trait values initially converged to the singular point, but then evolutionary branching occured, splitting the population into two distinct subpopulations (Fig. 2(e)).

Under the different-variance condition, the singular strategy is always convergence stable, but its evolutionary stability depends on the relative magnitude of male and female emergence variance. When the variance of male emergence timing σ_*M*_ is not small relative to that of female emergence timing σ_*F*_, the singular strategy is an ESS (Fig. 3(a) and Fig. 3(b)). When σ_*M*_ is sufficiently small relative to σ_*F*_, the singular strategy is no longer an ESS (Fig. 3(d)). The individual-based simulation showed that the trait values first converged to the singular point, then evolutionary branching was observed (Fig. 3(e)). It should be noted that the two subpopulations that emerged after branching were asymmetric in size: the early-emerging subpopulation (smaller trait values) was consistently smaller than the late-emerging subpopulation (larger trait values). This asymmetry is robust across different initial trait values and random seeds.

### Parameter dependence

To determine the generality of these results, we systematically analyzed how evolutionary outcomes depend on parameter values.

We first consider the singular strategy and examine the evolutionary and convergence stability of singular strategies under the neutral condition. Fig. S3 shows that the singular strategy consistently exhibits protandry. Fig. S4 and Fig. S5 show that, within the parameter values we used, the singular strategies are always convergence stable and ESS. Thus, evolutionary branching will not occur under this condition.

For the trade-off condition, we evaluated evolutionary stability by calculating the second derivative of the invasion fitness at the singular point, 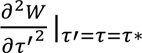, across a range of competitiveness function parameters: the inflection point *s*, the upper asymptote *h*, and the steepness *k* (Fig. 4). Blue regions satisfy condition (11), indicating a negative second derivative of the invasion fitness, which confirms that the singular point is an ESS. Conversely, red regions violate condition (11) (indicating a positive second derivative), meaning the singular point is not an ESS. We also confirmed that the singular points are convergence stable within these parameter sets (Fig. S6). Therefore, evolutionary branching occurs within the parameter set corresponding to the red regions. The evolutionarily unstable region (red region) expands as the parameter *k* increases. Furthermore, we observe that the singular strategy is not an ESS only in the intermediate range of *s*. In that range, the singular strategy is more likely to be not ESS as *h* increases. Fig. S7 shows the parameter dependence of the evolutionarily singular point. In all tested parameter sets, the singular points are negative. The value of the singular point decreases as *s* increases and *h* decreases. Furthermore, a lower mortality rate results in a more negative (smaller) singular point.

**Fig. 4.**
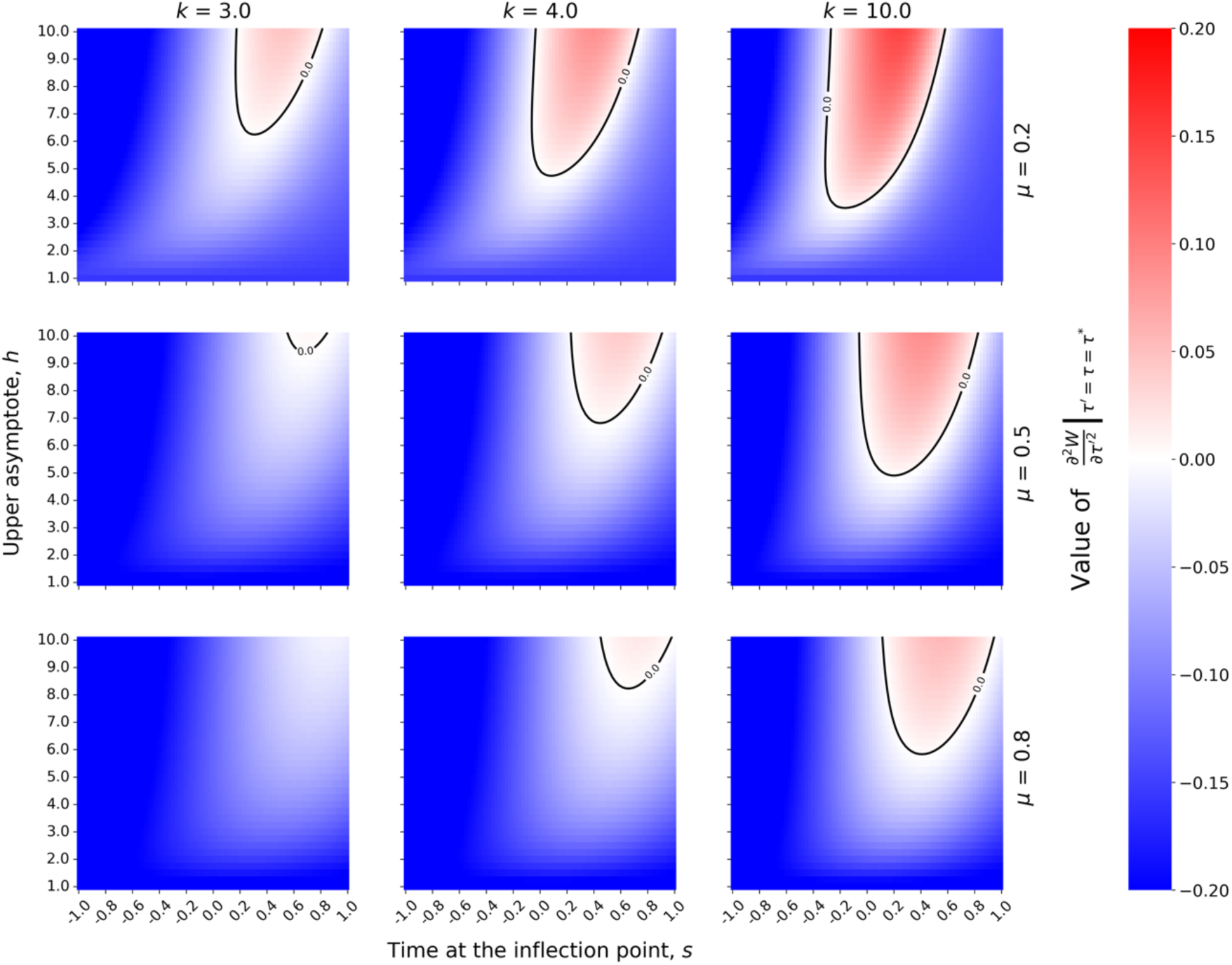
Evolutionarily stability analysis under the trade-off conditions. Heat maps show the second derivative of invasion fitness 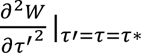 as a function of competitiveness parameters: inflection point (*s*), upper asymptote (*h*), steepness (*k*), and death rate (μ). Red regions indicate positive values where evolutionary branching can occur (singular strategy is not evolutionarily stable). Other parameters are σ_*M*_ = 1.0, and σ_*F*_ = 1.0.

For the different-variance condition, we examined how the second derivative of the invasion fitness, 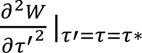, depends on the standard deviations of male (σ*_M_*) and female (σ_*F*_) emergence distributions (Fig. 5). All singular points are convergence stable in this parameter space (Fig. S8); therefore, as in Fig. 4, evolutionary branching occurs and polymorphism evolves in parameter regions, where the second derivative is positive (red regions). Evolutionary branching becomes more likely as the male emergence variance decreases relative to the female variance (i.e., the larger σ_*F*_ − σ_*M*_). Fig. S9 shows the parameter dependence of the evolutionarily singular point. Throughout the tested parameter range, the singular point remained negative. Similar to the trade-off condition, a lower mortality rate results in a more negative (smaller) singular point.

**Fig. 5.**
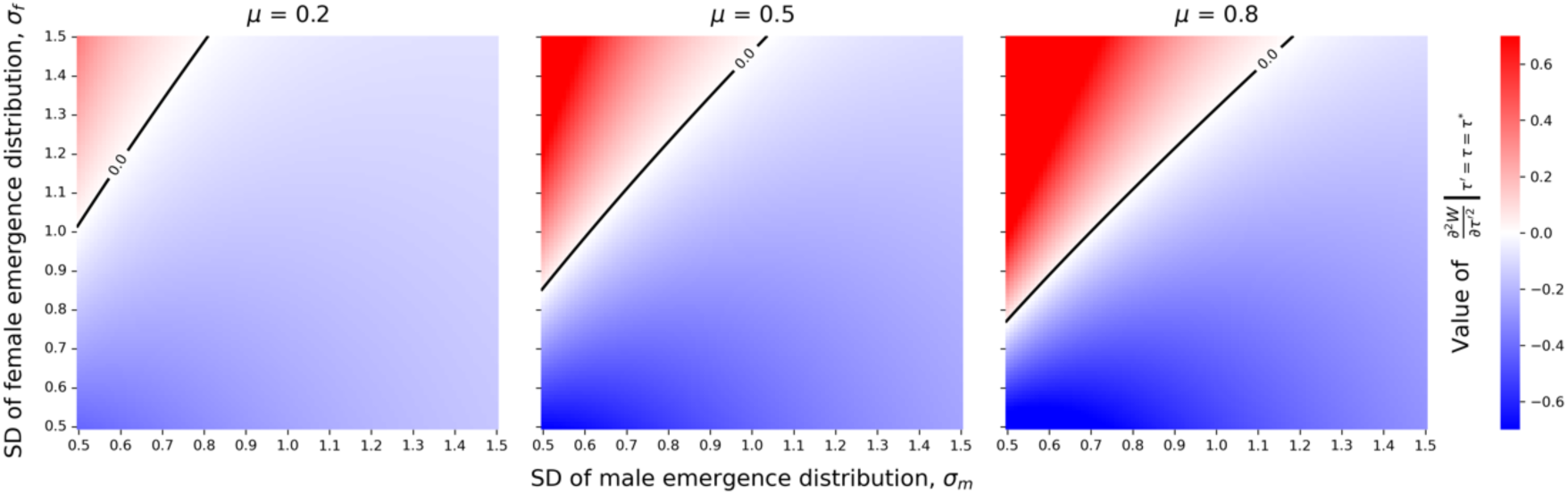
Effect of emergence distribution variances on evolutionarily stability. Heat map shows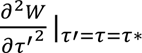 as a function of standard deviations of male (σ) and female (σ) emergence distribution, and death rate ( μ). Red regions indicate positive values of 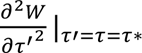, and thus conditions favoring evolutionary branching. Other parameters are *s* = 0.0, *h* = 1.0, and *k* = 1.0.

### Multiple branching under the different-variance condition

We found that multiple branching—the phenomenon where evolutionary branching occurs more than once, resulting in the emergence of three or more phenotypically discrete subpopulations—can occur under the different-variance condition (*c*(*t*) = 1 and σ_*M*_ ≠ σ_*F*_). Multiple branching tends to occur in the parameter space where the standard deviation in emergence timing of males (σ_*M*_) is sufficiently smaller than that of females (σ_*F*_). Fig. S10 shows the result from the individual-based simulation, illustrating the occurrence of multiple branching and the subsequent evolution of six distinct clusters. Fig. S11 shows the emergence probability and the number of individuals under such condition.

We investigate how parameter values influence the number of subpopulations that evolve through multiple branching. Fig. 6(a) shows the distributions of trait values in the final generation (generation 10000) of the individual-based simulations as a function of the ratio of standard deviations in emergence timing between males and females (σ_*m*_⁄σ_*f*_), when σ_*f*_ = 2.0. Monomorphic males evolve when σ_*M*_⁄σ_*F*_ > 0.7. However, as σ_*m*_⁄σ_*f*_ decreases, dimorphic males evolve, and as σ_*m*_⁄σ_*f*_ decreases further, the number of male subpopulations increases. Furthermore, almost all subpopulations have negative trait values, indicating that most clusters exhibit protandry.

**Fig. 6.**
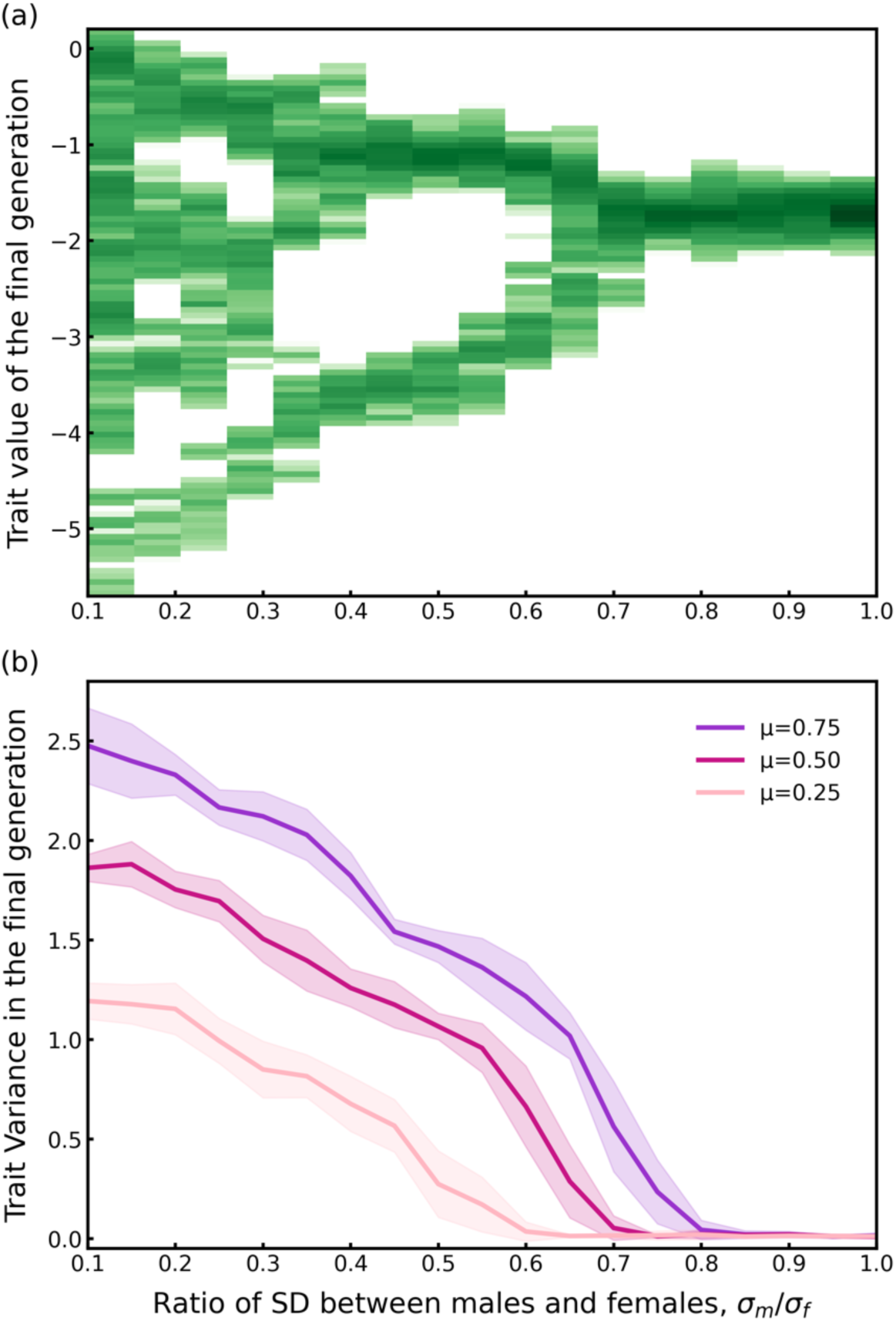
Parameter dependence of trait values in the final generation under multiple-branching conditions. (a) The distributions of trait values in the final generation of individual-based simulation are plotted as a function of the ratio of standard deviation in emergence timing between males and females (σ_*m*_⁄σ_*f*_), when σ_*f*_ = 2.0. Each parameter set was evaluated over 10 independent trials. Other parameters are μ = 0.5, *s* = 0.0, *h* = 1.0, *k* = 1.0, *N*_*M*_τ = 1000, *N*_*F*_ = 1000, *u* = 0.05, and σ_*mut*_ = 0.02. (b) The variance in trait values at generation 10000 of individual-based simulation are plotted as a function of σ_*m*_⁄σ_*f*_, when σ_*f*_ = 2.0. Each parameter set was evaluated over 10 independent trials. The solid lines represent the mean value, while the shaded regions indicate the standard deviation. Cases under three different death rates (μ = 0.25, 0.5, and 0.75) are shown. Other parameters are *s* = 0.0, *h* = 1.0, *k* = 1.0, *N*_*M*_τ = 1000, *N*_*F*_ = 1000, *u* = 0.05, and σ_*mut*_ = 0.02.

Fig. 6(b) shows the variance of trait values in the final generation as a function of σ_*m*_⁄σ_*f*_. The trait variance is close to zero in the parameter region where monomorphic males evolve. As σ_*m*_⁄σ_*f*_ decreases, the trait variance increases gradually. Furthermore, a higher death rate of adult males led to the evolution of polymorphism under weaker conditions, i.e., larger values of σ_*M*_⁄σ_*F*_. Even when comparing cases with the same σ_*m*_⁄σ_*f*_, higher death rate results in greater trait variance.

## Discussion

In this study, we demonstrated that male dimorphism in emergence timing can evolve as a consequence of disruptive selection arising from scramble competition for mating opportunities. Although several empirical studies have documented the coexistence of protandrous males and those emerging synchronously with females (Alcock, 1997; Kubo et al., 2025), classical models have consistently predicted unimodal protandry (Wiklund & Fagerström, 1977; Bulmer, 1983; Iwasa et al., 1983; Parker & Courtney, 1983). Our adaptive dynamics approach resolves this discrepancy by identifying two distinct ecological mechanisms that can generate evolutionary branching of male emergence timing.

### Two mechanisms for dimorphism evolution

The first mechanism operates through the trade-off between emergence timing and male competitiveness. Under this condition, dimorphism evolves when the relationship between development time and body size (and hence competitive ability) exhibits strong nonlinearity (i.e., a steep sigmoidal shape) (Fig. 2, Fig. 4). Disruptive selection then favors two distinct strategies: early-emerging males that maximize encounters with virgin females, and late-emerging males that invest in prolonged growth to achieve higher competitive ability. Previous protandry models that incorporated trade-offs assumed an exponentially increasing function (Zonneveld, 1996), which merely predicted delayed protandry rather than dimorphism. Our adoption of a sigmoidal function is biologically justified by the widespread occurrence of intraspecific variation in the number of larval instars observed in holometabolous insects, including butterflies and bees (Esperk et al., 2007). Because each additional molt results in a substantial and discrete increase in final body size (Nijhout, 1975), the transition in competitive ability is better captured by a sigmoidal rather than an exponential function.

The second mechanism operates through the asymmetry in emergence timing variance between the sexes. Under this condition, evolutionary branching occurs when the variance of the female emergence distribution (σ_*F*_) is sufficiently larger than that of males (σ_*M*_) (Fig. 3, Fig. 5). Notably, dimorphism evolves despite male competitiveness being independent of emergence timing. The underlying process can be interpreted as an analogy to evolutionary branching driven by resource competition (Doebeli & Dieckmann, 2000): virgin females represent a temporally distributed resource, and males are consumers competing for access to this resource. Selection initially favors an intermediate emergence timing where virgin females are abundant. However, once the population reaches the evolutionarily singular strategy, males with this intermediate timing experience intense competition, and phenotypes deviating from the singular strategy—both earlier and later—achieve higher fitness. This frequency-dependent selection drives evolutionary branching.

Furthermore, the plausibility of these mechanisms can be evaluated through empirical measurements in the field. In particular, simultaneous measurements of emergence timing, body size, and mating success in natural populations would allow us to test whether the assumed nonlinear relationship between development time and competitiveness is realized. In addition, quantifying the variance of emergence timing of each male and female would allow us to examine whether males and females differ in their emergence variability, as predicted by the different-variance mechanism. Thus, these empirical approaches provide a critical assessment of the evolutionary feasibility of these mechanisms.

### Biological implications of the two mechanisms

These two mechanisms generate distinct and empirically testable predictions. Under the trade-off mechanism, the two male morphs should differ substantially in body size, with late-emerging males being conspicuously larger. Under the variance asymmetry mechanism, body size differences between morphs should be minimal or absent, because competitiveness is decoupled from emergence timing. Existing empirical observations appear consistent with the trade-off mechanism: in both *Fabriciana nerippe* (Kubo et al., 2025) and *Amegilla dawsoni* (Alcock, 1997), late-emerging males exhibit markedly larger body sizes than early-emerging males. Whether systems conforming to the variance asymmetry mechanism exist in nature remains an open question worthy of empirical investigation.

Our parameter analyses revealed that the likelihood of evolutionary branching depends on specific combinations of biological parameters. Under the trade-off condition, branching is most likely when the competitiveness function has a high steepness (*k*) and an intermediate inflection point (*s*) (Fig. 4). Under the variance asymmetry condition, branching becomes increasingly likely as the ratio σ_*M*_/σ_*F*_decreases (Fig. 5). These parameter dependencies suggest that dimorphism should be more prevalent in species with pronounced developmental plasticity in body size and in species where male emergence is more synchronized than female emergence.

A key question is whether the condition σ_*F*_ > σ_*M*_ assumed in the variance asymmetry mechanism is biologically realistic. Although empirical studies directly comparing the variances of emergence timing between the sexes remain scarce, available evidence suggests that this condition may be common in protandrous insects. In the butterfly *Bicyclus anynana*, experiments revealed that the phenotypic variance of development time is larger in females than in males (Zwaan et al., 2008). More generally, because fecundity selection favors a larger female body size in many insect species, females tend to have longer larval development periods than males (Teder, 2014; Teder et al., 2021). Longer development times are often associated with greater developmental variance, as stochastic fluctuations accumulate over a longer growth period (Yurk & Powell, 2010). Together, these observations suggest a plausible mechanistic pathway that could generate the σ_*F*_ >σ_*M*_ condition required for evolutionary branching in our model. Testing this prediction systematically across species, for example by comparing sex-specific coefficients of variation in emergence timing, represents a promising e direction for future empirical research.

### Asymmetric morph frequencies

A notable prediction of our model is the asymmetry in subpopulation sizes following evolutionary branching. Under the different-variance condition, the early-emerging morph consistently comprises a smaller proportion of the population than the late-emerging morph (Fig. 3e). This asymmetry arises because the early-emerging niche—the period before peak female emergence—is temporally constrained compared to the broader window available to late-emerging males. Such frequency asymmetry between alternative morphs is commonly observed in natural populations and has typically been explained through condition-dependent threshold models (e.g., Hazel et al., 1990). Our results demonstrate that asymmetric morph frequencies can also emerge as a natural consequence of disruptive selection without invoking condition dependency.

### Multiple branching and the evolution of polymorphism

Another prediction of our model is the possibility of multiple branching events under the different-variance condition, leading to the evolution of three or more discrete phenotypic clusters (Fig. 6). Multiple branching occurs when σ_*M*_ is substantially smaller than σ_*F*_— that is, when male emergence is highly synchronized relative to female emergence. Under these conditions, even after an initial branching event, the resulting subpopulations may still experience disruptive selection, driving further diversification. This result extends beyond the typical two-strategy framework that dominates much of the alternative reproductive strategies literature (Gross, 1996; Oliveira et al., 2008) and suggests that the number of coexisting strategies may itself be an evolved outcome shaped by the temporal distribution of mating opportunities.

A higher adult male mortality (μ) facilitates the evolution of polymorphism under less stringent variance conditions (Fig. 6b). This counterintuitive result can be understood by considering that higher mortality reduces the temporal overlap among competing males, effectively partitioning the mating season into more distinct temporal niches. This prediction could be tested by comparing species or populations with different adult mortality rates.

### Model assumptions and limitations

Our model makes several simplifying assumptions that warrant discussion. First, we assumed that emergence timing is the sole trait under selection and that it evolves independently in males and females. In reality, genetic correlations between the sexes may constrain the evolution of sexual dimorphism in emergence timing (Lande, 1980). Second, we assumed that all females mate immediately upon emergence and mate only once. While this assumption is reasonable for many butterfly species, relaxing it to allow for female choice or multiple mating could substantially alter the selective landscape (Zonneveld, 1996). Third, our model does not incorporate stochastic environmental variation in emergence timing, which can influence the evolution of protandry through bet-hedging mechanisms (Iwasa & Haccou, 1994). A particularly important simplification is our treatment of the variances in emergence timing (σ_*M*_ and σ_*F*_) as fixed parameters rather than evolving traits. In our model, we analyzed how the probability distribution of male emergence timing evolves when only the mean emergence timing (τ) is allowed to evolve. However, the variance itself is likely subject to selection and may coevolve with the mean (Bruijning et al., 2020). Under the variance asymmetry mechanism, for example, selection might favor reduced variance in male emergence timing, which would in turn alter the conditions for evolutionary branching. Conversely, after branching occurs, selection might favor increased variance within each subpopulation to reduce competition among males with similar strategies. Extending the model to allow the simultaneous evolution of both the mean and variance of emergence timing—potentially using function-valued trait approaches (Dieckmann et al., 2006) or quantitative genetic models such as oligomorphic dynamics (Sasaki & Dieckmann, 2011)—represents a promising direction for future work. Such an extension could reveal whether the variance asymmetry required for branching can itself evolve, and whether the observed dimorphism in natural populations represents a stable evolutionary outcome or a transient state.

Finally, our model assumes that emergence timing is a fixed genetic trait, and that dimorphism arises as a genetic polymorphism. In nature, however, alternative reproductive phenotypes are often condition-dependent, where individuals adopt a tactic based on their physiological status or environmental conditions (Gross, 1996). For instance, in the solitary bee *A. dawsoni*, some studies suggest that the coexistence of early-and late-emerging males represents a genetically fixed mixed evolutionarily stable strategy (Alcock, 1997), whereas others propose a condition-dependent tactic (Beveridge et al., 2006). Extending our framework to incorporate the condition-dependent tactic expression represents a promising direction for future work.

### Concluding remarks

The framework developed here applies broadly to any system where reproductive success depends on the temporal overlap between the sexes, including fishes, amphibians, reptiles (Morbey & Ydenberg, 2001), migratory birds (Kokko, 1999), and flowering plants (Augspurger, 1981; Forrest, 2014). Our results contribute to the theoretical understanding of how continuous phenotypic variation can evolve into discrete polymorphisms through frequency-dependent selection. Notably, the evolutionary branching process we describe—whereby a unimodal population diverges into distinct phenotypic clusters— parallels the early stages of speciation driven by disruptive selection (Dieckmann & Doebeli, 1999; Yamaguchi et al. 2026). If assortative mating were to evolve between early- and late-emerging males, the intrapopulation polymorphism documented here could provide a substrate for incipient reproductive isolation. A plausible pathway is as follows: because early- and late-emerging males mate predominantly with females that emerge at different times within the season, and if female emergence timing is at least partly heritable, daughters of early males would tend to emerge earlier and daughters of late males later. Over generations, this temporal assortment of mating pairs could generate a genetic correlation between male strategy and female emergence timing, effectively producing allochronic reproductive isolation without requiring the evolution of mate-choice mechanisms *per se* (Hendry & Day, 2005). Such a scenario is analogous to “magic trait” models of speciation, in which the trait under disruptive selection simultaneously generates assortative mating as a byproduct (Servedio et al., 2011). Whether this pathway is realized in nature depends on the heritability of female emergence timing, the strength of temporal segregation between morphs, and the extent of genetic correlation between male and female emergence schedules. In this sense, the evolution of alternative reproductive strategies in timing may represent not only a stable endpoint but also an initial step toward species diversification.

