## Appendix for "Evolutionary branching of male emergence timing: Trade-offs and variance asymmetry as drivers of dimorphism"

### Appendix1

**Derivation of $\boldsymbol{C(t|\tau)}$**

Here, we derive the transformation from Eq. (2) to Eq. (3) in the main text. Starting from Eq. (2), we have:

$$\frac{dC(t|\tau)}{dt} + \mu C(t|\tau) = f(t)P_M(t|\tau).$$

Multiply both sides by $\exp[\mu t]$:

$$\frac{dC(t|\tau)}{dt}\exp[\mu t] + \mu C(t|\tau)\exp[\mu t] = f(t)P_M(t|\tau)\exp[\mu t].$$

Applying the product rule, we obtain:

$$\frac{d}{dt}\{\exp[\mu t]C(t|\tau)\} = f(t)P_M(t|\tau)\exp[\mu t].$$

Integrate both sides from $-\infty$ to $t$:

$$[\exp[\mu t]C(t|\tau)]_{-\infty}^{t} = \int_{-\infty}^{t} f(t_e)P_M(t_e|\tau)\exp[\mu t_e]\,dt_e,$$

$$\exp[\mu t]\,C(t|\tau) - \lim_{s\to-\infty}\{\exp[\mu s]\,C(s|\tau)\} = \int_{-\infty}^{t} f(t_e)P_M(t_e|\tau)\exp[\mu t_e]\,dt_e,.$$

Assuming $C(t|\tau) \to 0$ as $t \to -\infty$,

$$C(t|\tau) = \int_{-\infty}^{t} f(t_e)P_M(t_e|\tau)exp[-\mu(t-t_e)]dt_e.$$

### Appendix2

**Derivation of $W(\tau'|\tau)$**

Here, we derive the transformation from Eq. (6) to Eq. (7) in the main text. Starting from Eq. (6), we have:

$$W(\tau'|\tau) = \int_{-\infty}^{\infty} P_M(t_e|\tau')\,\phi(t_e|\tau,\tau')dt_e$$

$$= \int_{-\infty}^{\infty} P_M(t_e|\tau')\int_{t_e}^{\infty} M(t_e,t|\tau,\tau')\exp\,[-\mu(t-t_e)]dt\,dt_e$$

$$= \int_{-\infty}^{\infty} P_M(t_e|\tau')\int_{t_e}^{\infty} f(t_e)\frac{N_F P_F(t)}{N_{M_\tau}C(t|\tau)+N_{M_{\tau'}}C(t|\tau')}\exp\,[-\mu(t-t_e)]dt\,dt_e.$$

Taking the limit as the number of mutant individuals $N_{M_{\tau'}} \to 0$ and assuming $N_{M_\tau} = N_F$:

$$W(\tau'|\tau) = \int_{-\infty}^{\infty} P_M(t_e|\tau')\int_{z}^{\infty} f(t_e)\frac{P_F(t)}{C(t|\tau)}\exp\,[-\mu(t-t_e)]dt\,dt_e,$$

$$W(\tau'|\tau) = \int_{-\infty}^{\infty}\int_{t_e}^{\infty} P_M(t_e|\tau')f(t_e)\frac{P_F(t)}{C(t|\tau)}\exp\,[-\mu(t-t_e)]dt\,dt_e.$$

By changing the order of integration, we get

$$= \int_{-\infty}^{\infty}\int_{-\infty}^{t} P_M(t_e|\tau')\frac{f(t_e)}{C(t|\tau)}P_F(t)\exp\,[-\mu(t-t_e)]dt_e\,dt$$

$$= \int_{-\infty}^{\infty}\frac{P_F(t)}{C(t|\tau)}\int_{-\infty}^{t} f(t_e)P_M(t_e|\tau')\exp\,[-\mu(t-t_e)]dt_e\,dt$$

$$= \int_{-\infty}^{\infty}\frac{C(t|\tau')}{C(t|\tau)}P_F(t)\,dt.$$

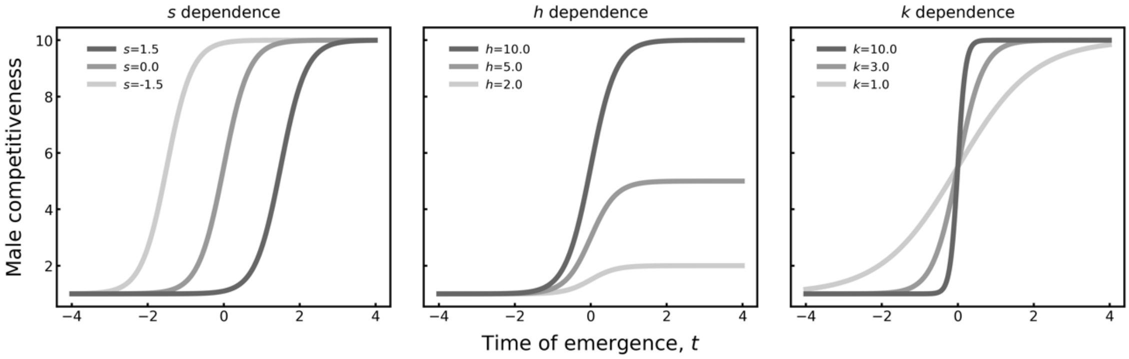


**Fig. S1** The outline of male competitiveness function. This figure illustrates how the male competitiveness function changes when varying one parameter at a time while keeping the others fixed at their default values ($s = 0.0$, $h = 10.0$, and $k = 3.0$).

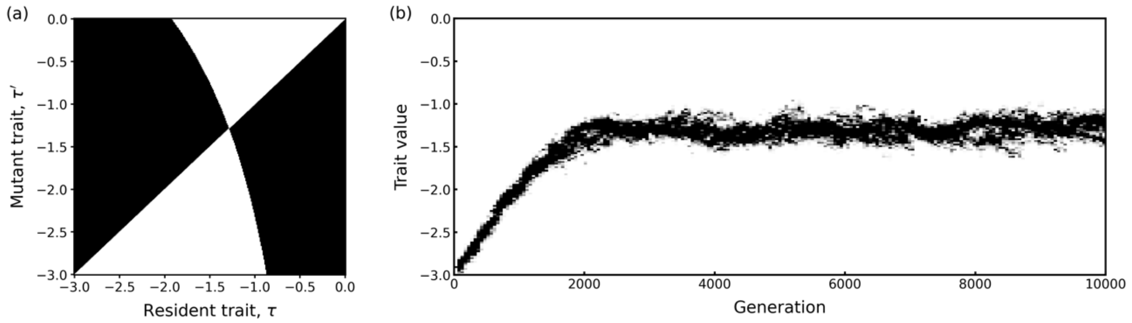


**Fig. S2** Pairwise invasibility plot (PIP) and evolutionary dynamics generated by the individual-based simulation under the neutral condition. (a) PIPs showing invasion fitness $W(\tau'|\tau)$ where $\tau$ and $\tau'$ represent resident and mutant trait values, respectively. Black regions indicate $W(\tau'|\tau) > 1$, where mutants can invade the resident population. (b) Individual-based simulation results showing trait distribution over 10000 generations. Parameters are $\sigma_M = 1.0$, $\sigma_F = 1.0$, $\mu = 0.5$, $s = 0.0$, $h = 1.0$, $k = 1.0$, $N_{M_\tau} = 1000$, $N_F = 1000$, $u = 0.05$, and $\sigma_{mut} = 0.02$.

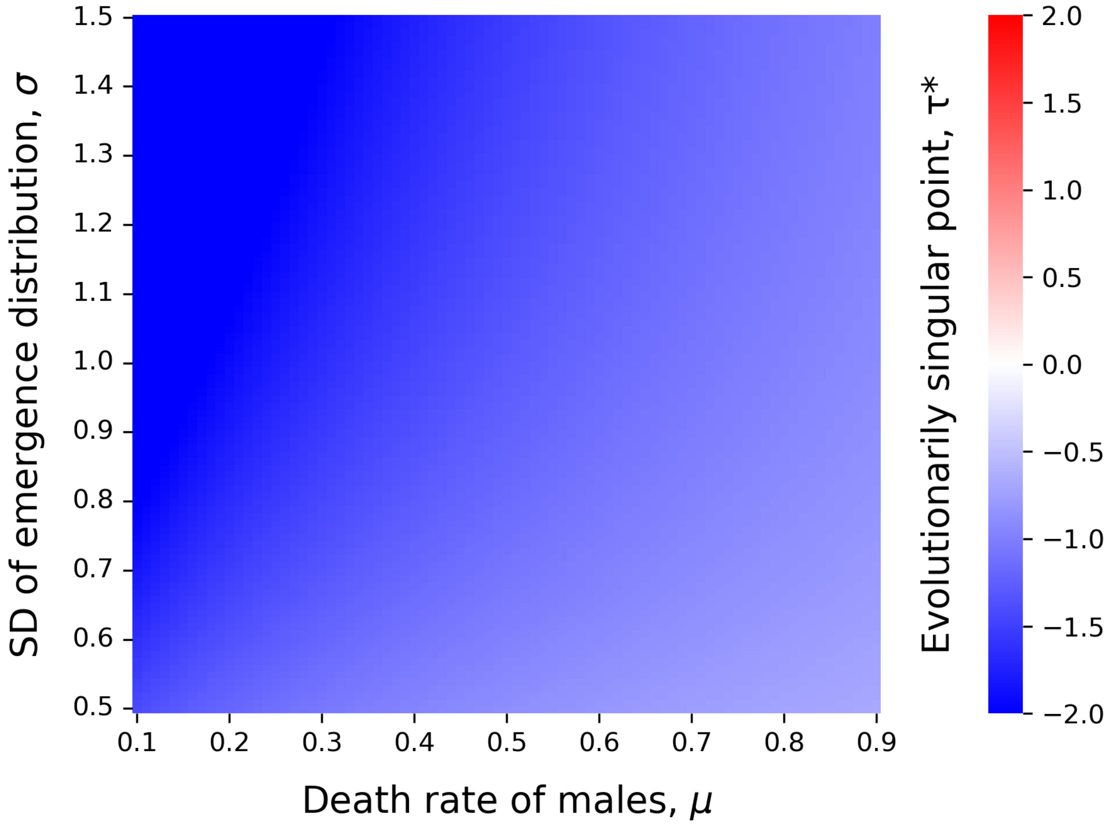


**Fig. S3** Parameter dependence of evolutionarily singular point under the neutral conditions. Heat map shows the evolutionarily singular point $\tau*$ as a function of parameters: Death rate of adult males (μ), standard deviations of emergence distribution ($\sigma = \sigma_M = \sigma_F$). Blue regions indicate negative values where evolutionarily singular strategy is ESS. Other parameters are $s = 0.0$, $h = 1.0$, and $k = 1.0$.

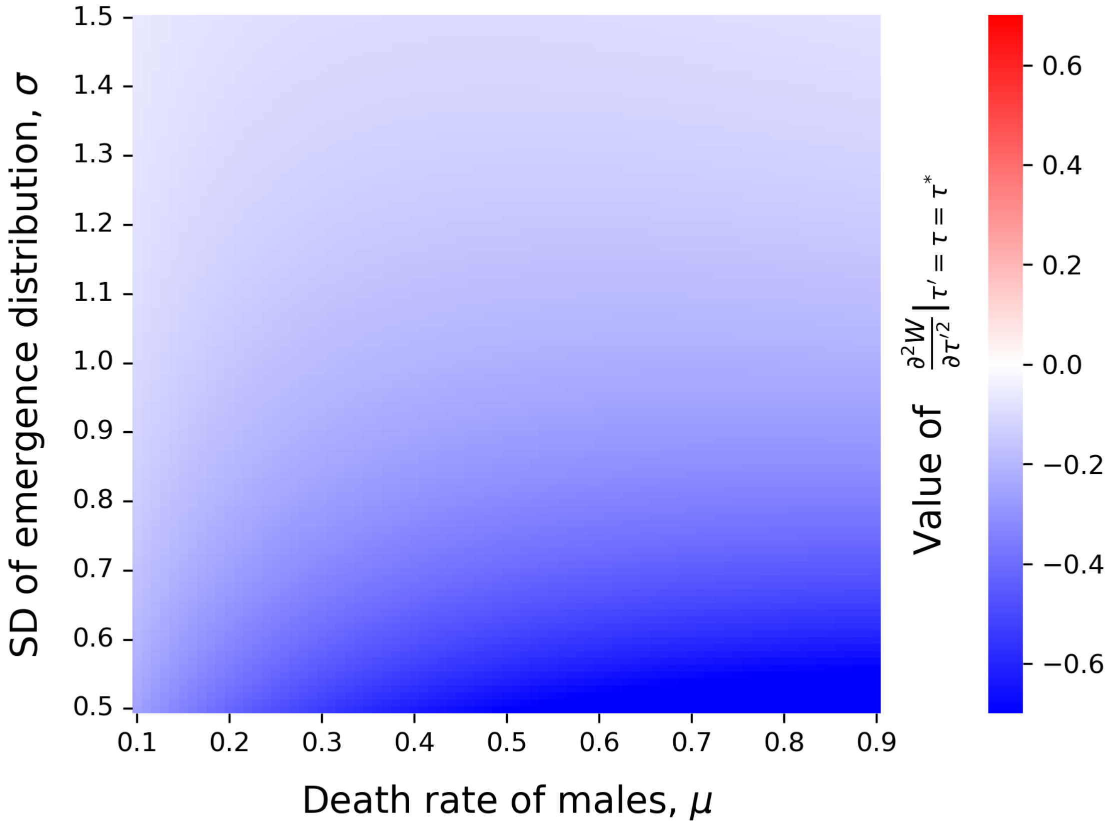


**Fig. S4** Evolutionarily stability analysis under the neutral conditions. Heat map shows the second derivative on invasion fitness $\frac{\partial^2 W}{\partial \tau'^2}|_{\tau'=\tau=\tau*}$ as a function of parameters: Death rate of adult males ($\mu$), standard deviations of emergence distribution ($\sigma = \sigma_M = \sigma_F$). Blue regions indicate negative values where evolutionarily singular strategy is ESS. Other parameters are $s = 0.0$, $h = 1.0$, and $k = 1.0$.

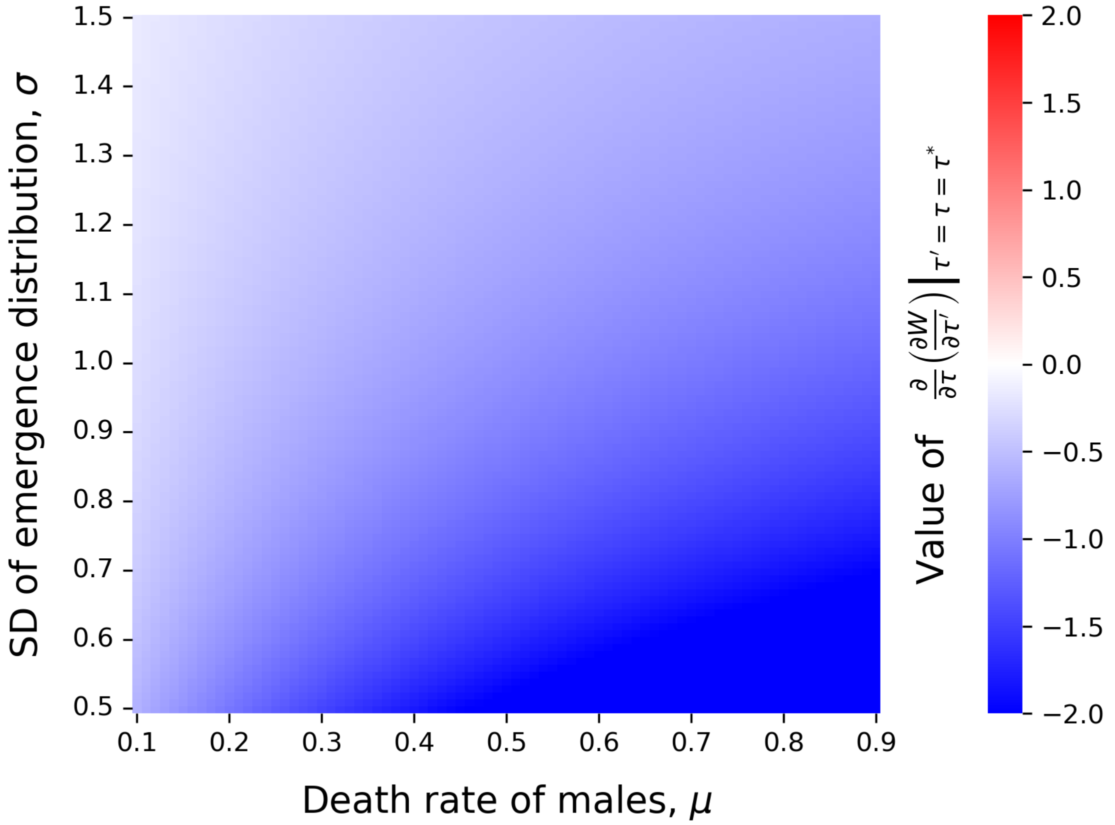


**Fig. S5** Convergence stability analysis under the neutral conditions. Heat map shows $\frac{\partial}{\partial\tau}(\frac{\partial W}{\partial\tau'})|_{\tau'=\tau=\tau*}$ as a function of parameters: Death rate of adult males ($\mu$), standard deviations of emergence distribution ($\sigma = \sigma_M = \sigma_F$). Blue regions indicate negative values where evolutionarily singular strategy is convergence stable. Other parameters are $s = 0.0$, $h = 1.0$, and $k = 1.0$.

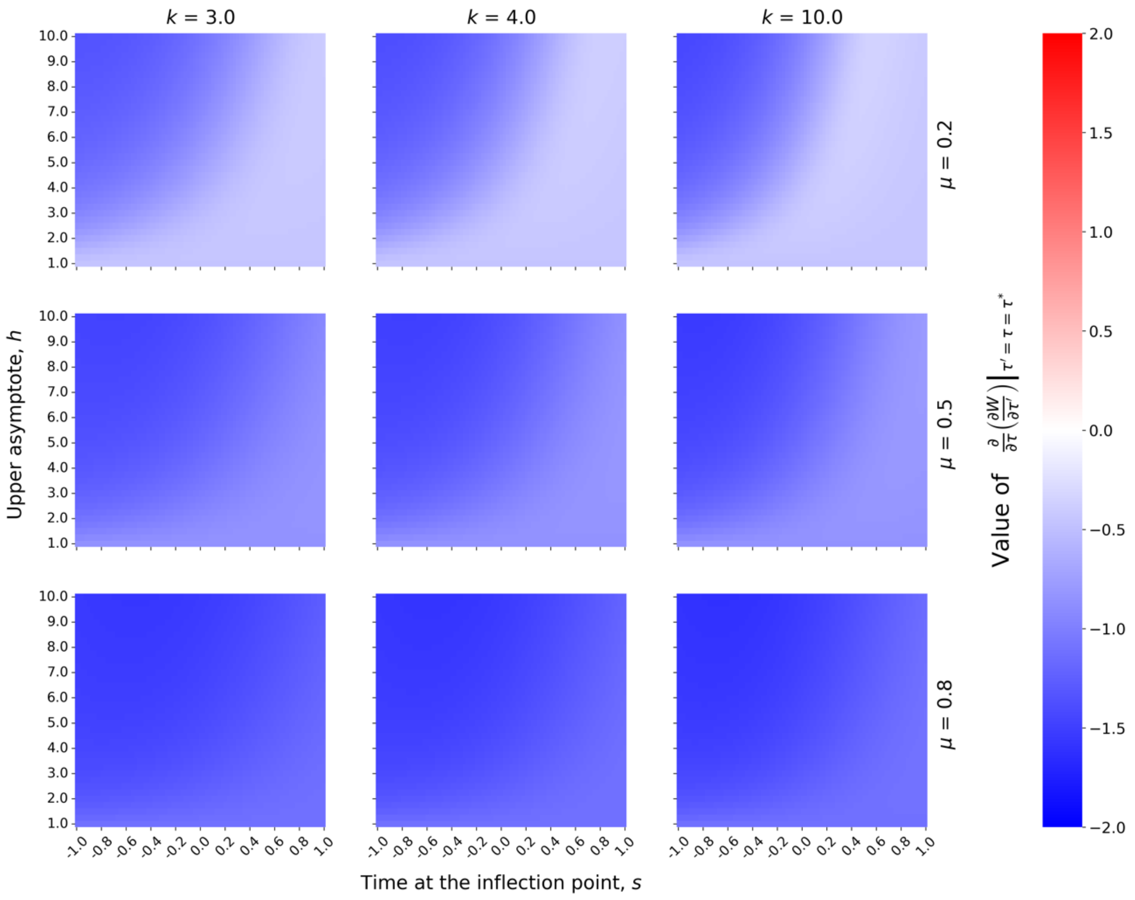


**Fig. S6** Convergence stability analysis under the trade-off conditions. Heat maps show $\frac{\partial}{\partial \tau}(\frac{\partial W}{\partial \tau'})|_{\tau'=\tau=\tau*}$ as a function of competitiveness parameters: inflection point ($s$), upper asymptote ($h$), and steepness ($k$). Blue regions indicate negative values where evolutionarily singular strategy is convergence stable. Other parameters are $\sigma_M = 1.0$, $\sigma_F = 1.0$, and $\mu = 0.5$.

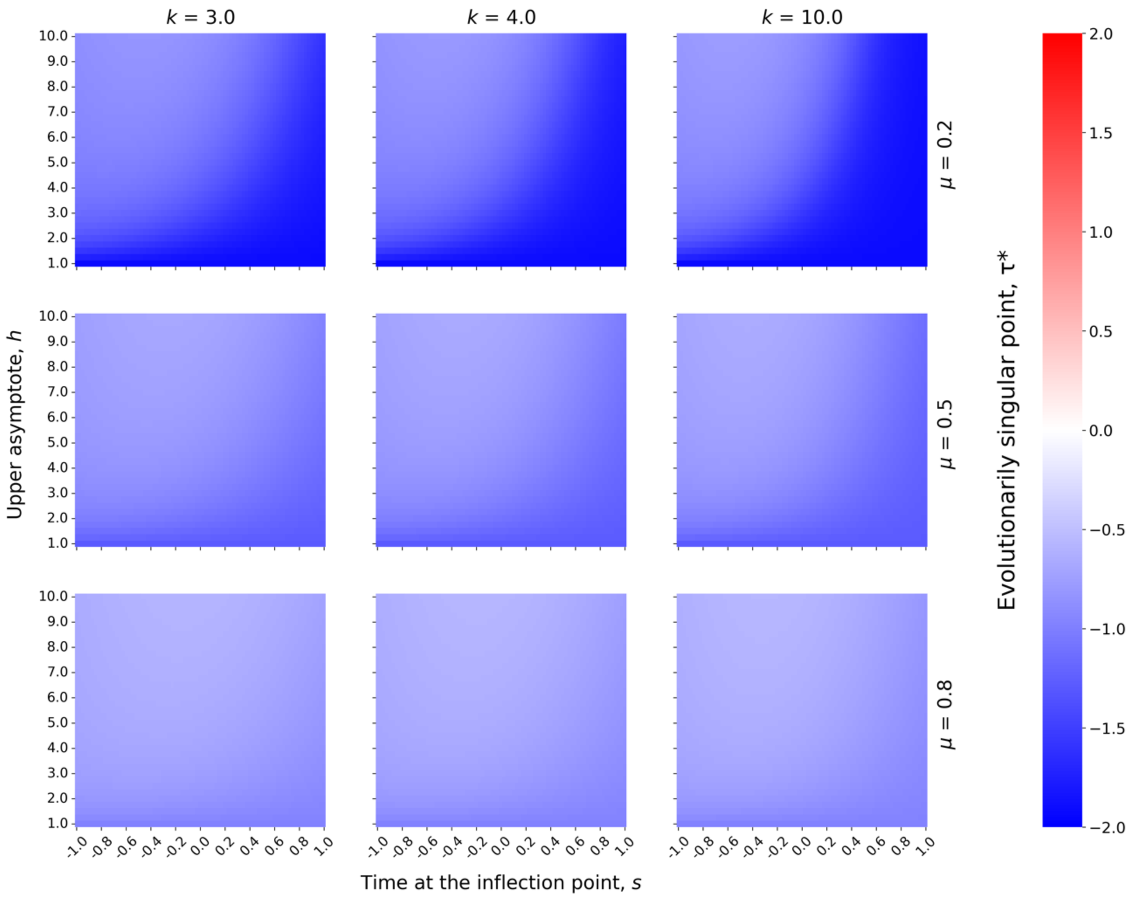


**Fig. S7** Parameter dependence of evolutionarily singular point under the trade-off conditions. Heat map shows the evolutionarily singular point $\tau*$ as a function of parameters: inflection point ($s$), upper asymptote ($h$), and steepness ($k$). Blue regions indicate negative values where evolutionarily singular strategy is convergence stable. Other parameters are $\sigma_M = 1.0$, $\sigma_F = 1.0$, and $\mu = 0.5$.

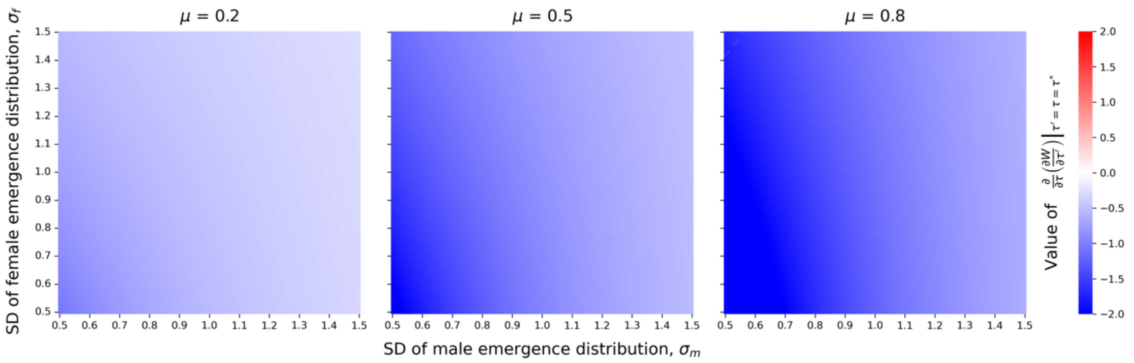


**Fig. S8** Convergence stability analysis under the different-variance conditions. Heat maps show $\frac{\partial}{\partial\tau}(\frac{\partial W}{\partial\tau'})|_{\tau'=\tau=\tau*}$ as a function of standard deviations of male ($\sigma_M$) and female ($\sigma_F$) emergence distribution. Blue regions indicate negative values where evolutionarily singular strategy is convergence stable. Other parameters are $\mu = 0.5$, $s = 0.0$, $h = 1.0$, and $k = 1.0$.

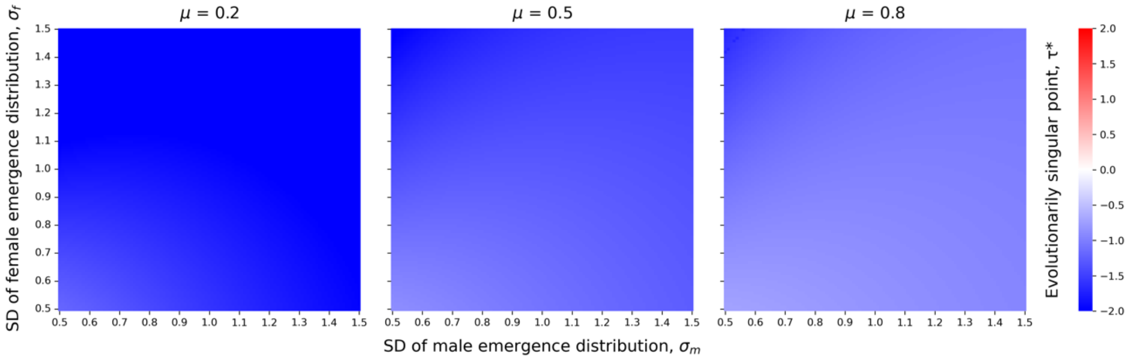


**Fig. S9** Parameter dependence of evolutionarily singular point different-variance conditions. Heat map shows the evolutionarily singular point $\tau^*$ as a function of standard deviations of male ($\sigma_M$) and female ($\sigma_F$) emergence distribution. Blue regions indicate negative values where evolutionarily singular strategy is convergence stable. Other parameters are $\mu = 0.5$, $s = 0.0$, $h = 1.0$, and $k = 1.0$.

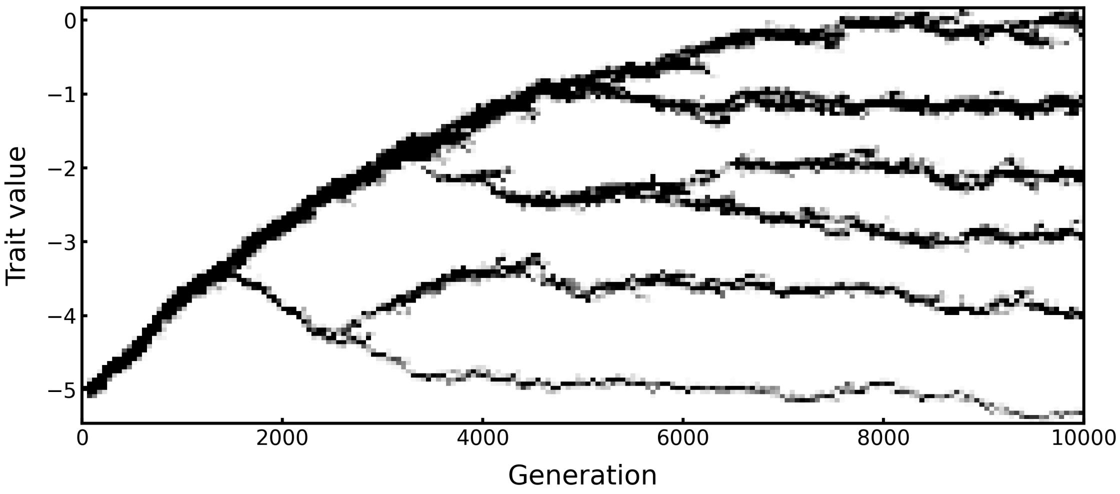


**Fig. S10** The evolutionary dynamics generated by the individual-based simulation, showing multi-branching. Result shows trait distribution over 10000 generations. Parameters are $\sigma_M = 0.25$, $\sigma_F = 2.0$, $\mu = 0.5$, $s = 0.0$, $h = 1.0$, $k = 1.0$, $N_{M_\tau} = 1000$, $N_F = 1000$, $u = 0.05$, and $\sigma_{mut} = 0.02$.

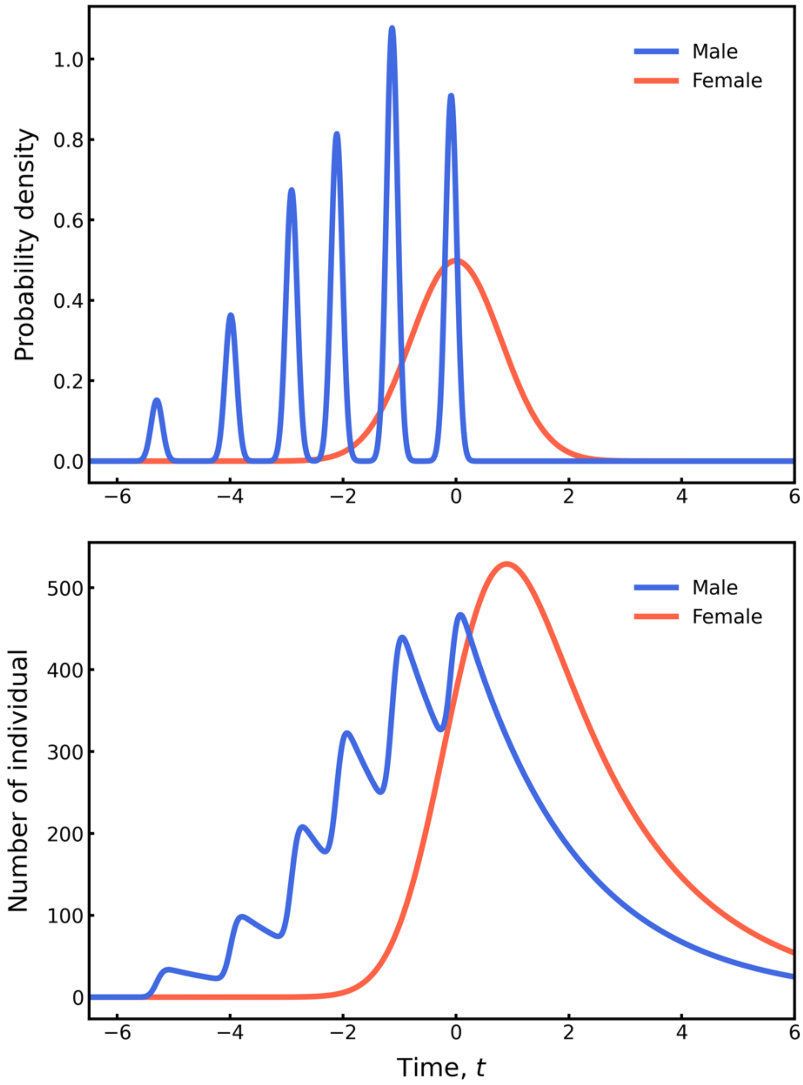


**Fig. S11** The emergence probability (top) and number of individuals (bottom) under the multi-branching scenario. The top panel is constructed as the weighted sum of normal distributions, where each distribution has its own mean and frequency obtained from the results of the individual-based simulations, while sharing a common variance. The bottom panel is calculated using equation (3), taking the top panel as the emergence probability, $P_M$, and then multiplying by the total number of males, $N_M$. Parameters are $\sigma_M = 0.25$, $\sigma_F = 2.0$, $\mu = 0.5$, $s = 0.0$, $h = 1.0$, $k = 1.0$, $N_{M_\tau} = 1000$, $N_F = 1000$, $u = 0.05$, and $\sigma_{mut} = 0.02$.
